# Cross-Genera Amplification of *prs/hlyA* by Multiplex PCR Resulted in Misidentification of *Enterococcus faecium* as *Listeria monocytogenes*

**DOI:** 10.64898/2026.01.20.700535

**Authors:** H.B. Ali, A.S. Kumurya, J.M. Ajagbe, Yahaya Usman, Abubakar Sadiq Baba, M. Usman

**Author notes:** Corresponding Author (H.B. Ali).

## Abstract

**Importance:** Molecular confirmation of *Listeria monocytogenes* typically employs a multiplex PCR method that targets both genus-specific prs and species-specific hlyA genes. This study assessed the specificity of this assay within Nigeria, where local microbial diversity may influence performance outcomes.

**Methods:** Out of eight phenotypically presumptive *L. monocytogenes* food isolates tested, six produced the expected prs and hlyA amplicons, with five classified as serogroup 1/2b. However, all six isolates tested negative for the crucial virulence regulator *prfA*, necessitating further investigation.

**Results:** Definitive 16S rRNA gene sequencing revealed that only two of the six PCR-positive isolates were identified as L. monocytogenes, while the remaining four were identified as *Enterococcus faecium*. This results in a false-positive rate of 66.7% (4/6) for the assay in this particular context. Phylogenetic analysis corroborated the taxonomic distinction, exhibiting a robust clustering of the four *E. faecium* isolates with reference strains. In contrast, the two confirmed *L. monocytogenes* isolates formed a separate sub-clade, indicating regional divergence and further underscoring the assay’s inability to differentiate between *L. monocytogenes* and Enterococcus species.

**Conclusion:** These findings highlight a significant lack of specificity, as the *prs/hlyA* primers exhibited cross-reactivity with non-target *E. faecium*. The anomalous negative result for *prfA* served as a critical diagnostic indicator. Consequently, the positive outcomes from this widely utilized confirmatory assay should be regarded as presumptive and necessitate additional verification.

## 1.0 INTRODUCTION

Prompt and precise identification of *Listeria* monocytogenes, a significant foodborne pathogen that leads to invasive listeriosis, is crucial for clinical diagnostics and food safety monitoring. Although traditional culture-based methods remain important, molecular techniques offer essential speed and sensitivity for verification. Among these, the multiplex polymerase chain reaction (PCR) assay created by Doumith and colleagues—which simultaneously targets the genus-specific prs gene and the L. monocytogenes-specific hlyA gene (which encodes listeriolysin O)—has become a commonly used standard for distinguishing *L. monocytogenes* from other *Listeria* species in both clinical settings and food samples [1, 2].

The dependability of a diagnostic assay largely depends on its specificity within the microbial environment in which it is utilized. Primer sequences that have been validated against certain strain collections might not consider the genetic variability present in various geographical or environmental settings, which could result in cross-reactivity and false-positive findings. [3]. This is especially important in areas where the local microbial phylogeny is not well defined and where closely related Gram-positive bacteria (such as Enterococcus spp. and Brochothrix spp.) might have conserved genomic regions in common. In an earlier surveillance study conducted in northeastern Nigeria, we isolated eight strains of *L. monocytogenes* that were phenotypically presumed from various food samples using standard biochemical methods and automated systems (VITEK 2)[4]. This created an opportunity to genetically analyze these isolates while simultaneously assessing the effectiveness of the Doumith multiplex PCR in this less-explored context. The main aim of this study was not to conduct surveillance, but rather to perform a critical diagnostic evaluation: to determine the agreement between this established PCR method and definitive 16S rRNA gene sequencing for isolates from a region with unique microbial ecology.

The study uncovered a significant and surprising lack of agreement, resulting in the identification of considerable cross-genera amplification. This report outlines the assay’s specificity failure, where *Enterococcus faecium* was wrongly identified as L. monocytogenes, and discusses the immediate ramifications for diagnostic practices that depend on this widely used molecular confirmation test.

## 2.0 MATERIALS AND METHODS

### 2.1. Study Design and Bacterial Isolates

This diagnostic evaluation study employed eight isolates of *Listeria monocytogenes* that were presumptively identified based on their phenotypic characteristics (labeled BO1–BO8), which were sourced from a previous food surveillance investigation conducted in Maiduguri, Nigeria. [5]. The main goal was to assess the specificity of a standard multiplex PCR assay in comparison to definitive 16S rRNA gene sequencing. Isolates were taken from −80°C storage in Brain Heart Infusion broth containing 20% glycerol, then subcultured on Blood Agar to ensure purity and incubated at 37°C for 24 hours before analysis. A summary of the sample processing workflow from the initial surveillance study can be found in Supplementary Table S1.

### 2.2. DNA Extraction

Genomic DNA was obtained from fresh, pure colonies utilizing the Quick-DNA™ Fungal/Bacterial Microprep Kit (Zymo Research, USA), following the manufacturer’s instructions for bacterial cells. The concentration and purity of the DNA were assessed using a NanoDrop™ One spectrophotometer (Thermo Fisher Scientific). The extracted DNA was then stored at −20°C.

### 2.3. Multiplex PCR Assay under Evaluation

The effectiveness of the commonly utilized Doumith *prs/hlyA* multiplex PCR test [3] was evaluated. All oligonucleotide primers utilized in this research are detailed in supplementary Table S2.

#### 2.3.1. Confirmation of Genus and Species (prs and hlyA)

The multiplex reaction was conducted in a 25 µL volume that included: 12.5 µL of 2× OneTaq Quick-Load Master Mix (New England Biolabs), 0.5 µM of each primer, 5 µL of template DNA, along with nuclease-free water. The thermocycling conditions were: an initial denaturation step at 95°C for 5 minutes; followed by 35 cycles of 95°C for 30 seconds, 58°C for 30 seconds, 72°C for 45 seconds; and a final extension at 72°C for 5 minutes.

#### 2.3.2. Detection of Serogroup and Virulence Genes

A distinct multiplex PCR was conducted for further characterization of isolates and to examine the *prfA* virulence regulator, while serogrouping was accomplished using primers for serogroups 1/2a, 1/2b, and 4b [3, 6]. The composition of the reaction was the same as described in Section 2.3.1. The cycling conditions included: 94°C for 5 minutes; followed by 35 cycles of 94°C for 40 seconds, 53°C for 75 seconds, and 72°C for 75 seconds; concluding with 72°C for 7 minutes.

### 2.4 Confirmatory 16S rRNA Gene Sequencing and Phylogenetic Analysis

#### 2.4.1. Amplification and Sequencing

The near-complete 16S rRNA gene was amplified using the universal primers 27F and 1492R [7] In a 50 µL reaction, the setup included 25 µL of 2× Master Mix, 0.3 µM of each primer, and 5 µL of DNA. The cycling conditions were set at 95°C for 5 minutes; followed by 35 cycles of 95°C for 30 seconds, 55°C for 45 seconds, and 72°C for 1 minute, with a final extension at 72°C for 10 minutes. Amplicons were purified using the QIAquick PCR Purification Kit from Qiagen, and sequencing was performed bidirectionally using Sanger chemistry on an ABI 3500xl Genetic Analyzer from INQABA Biotechnical Industries in Nigeria.

For sequence analysis and phylogenetics, forward and reverse reads were assembled and edited using BioEdit version 7.2 [8]. Consensus sequences were matched against the NCBI GenBank database through BLASTn. For the purpose of phylogenetic analysis, sequences were aligned with reference strains utilizing ClustalW within MEGA11 [9]. The evolutionary lineage was deduced using the Neighbor-Joining approach [10] with 1,000 bootstrap iterations [11]. Evolutionary distances were calculated using the Maximum Composite Likelihood technique [12]. All uncertain positions were discarded (pairwise deletion option).

### 2.5. Gel Electrophoresis

PCR products (8 µL) were combined with 2 µL of 6× loading dye and subjected to separation on a 1.5% agarose gel in 0.5× TBE buffer, which was stained with Midori Green Advanced DNA Stain. The electrophoresis was carried out at 115 V for 25 minutes. A 100 bp DNA ladder was included to aid in size estimation. The gels were examined using a Bio-Rad Gel Doc™ XR+ system.

### 2.6. Data and Performance Analysis

The false-positive rate for the *prs/hlyA* assay was determined using the formula:

(Number of isolates that tested PCR-positive but were identified as non-*L. monocytogenes* through sequencing) / (Total number of PCR-positive isolates) × 100%.

Sequencing-based identification was deemed conclusive. The 16S rRNA gene sequences obtained in this study have been submitted to GenBank with the accession numbers PV809878.1 to PV809883.1.

## 3.0 RESULTS

### 3.1. The *prs/hlyA* Multiplex PCR Assay’s performance

Based on the simultaneous amplification of the anticipated 370 bp and 456 bp fragments, respectively, the multiplex PCR assay targeting the prs and hlyA genes identified six of the eight presumed isolates (BO1–BO6) as *Listeria monocytogenes* (Fig 1A). Since isolates BO7 and BO8 tested negative for both targets, they were not included in the analysis. A significant abnormality was noted, though: in the supplemental multiplex reaction, none of the six PCR-positive isolates generated an amplicon for the *prfA* virulence regulator gene (1060 bp).

**Figure 1.**
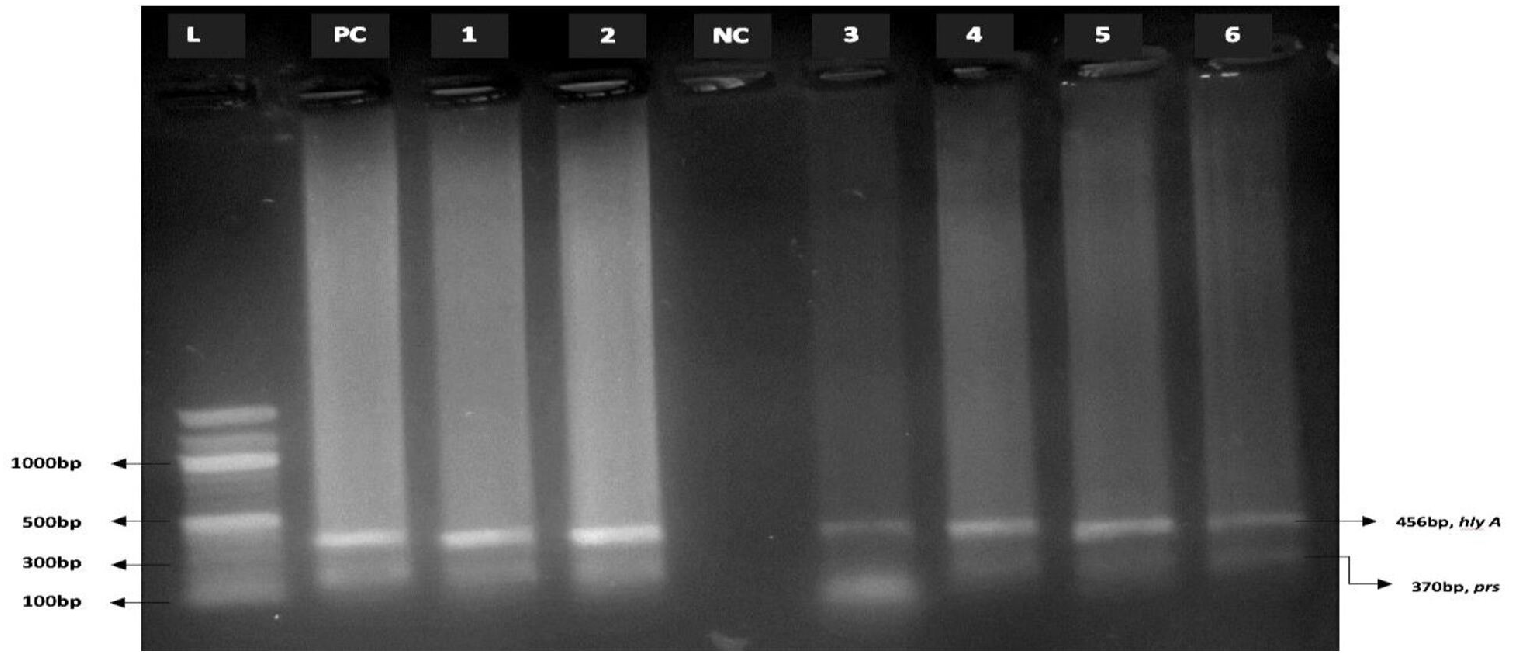
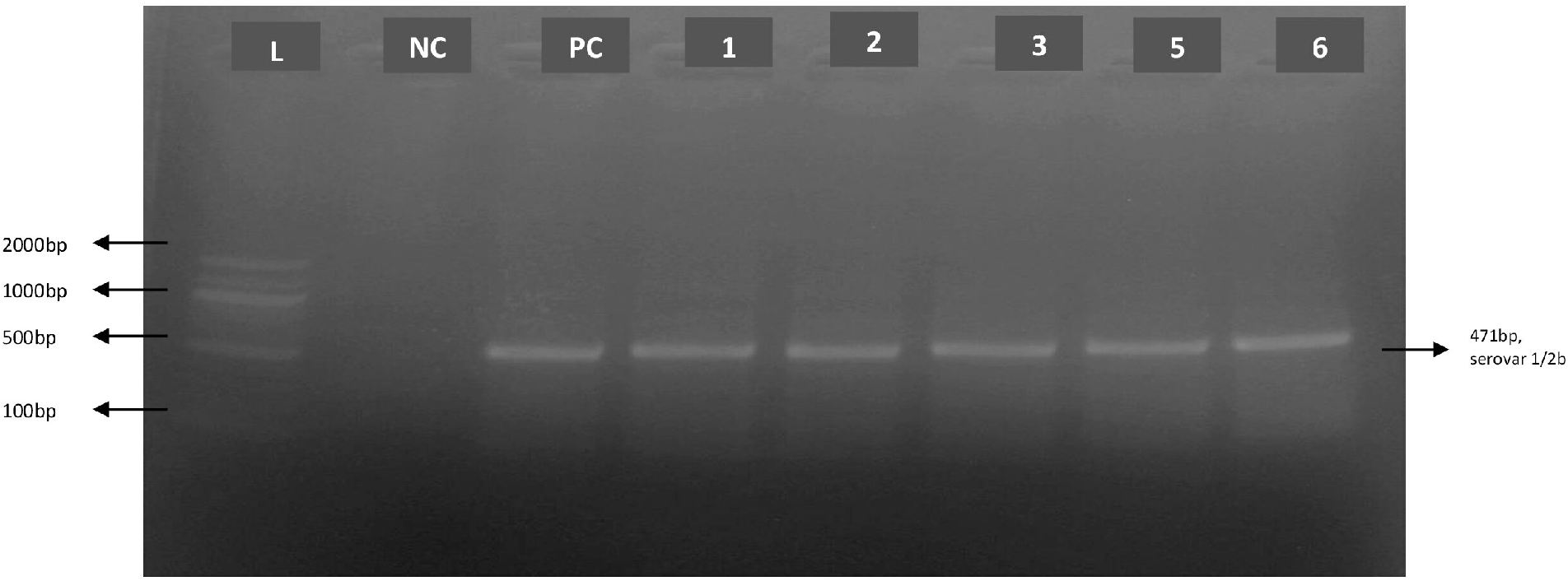
Assessment of the *prs/hlyA* multiplex PCR assay. (A) Gel electrophoresis used for identifying the *Listeria* genus (prs, 370 bp) and the *L. monocytogenes* species (hlyA, 456 bp). Lane M: 100 bp DNA marker; Lane PC: Positive control (*L. monocytogenes* ATCC 19115); Lane NC: Negative control (nuclease-free water); Lanes 1-6: Isolates BO1 to BO6. (B) Serogroup analysis for lineages 1/2a (691 bp), 1/2b (471 bp), and 4b (597 bp). Lane identifications are the same as in (A). A 471 bp band (serogroup 1/2b) is detected in lanes 1, 2, 3, 4, and 6.

### 3.2. 16S rRNA Gene Definitive Identification Sequencing Exposes Important Disagreement

All six PCR-positive isolates underwent 16S rRNA gene sequencing in order to address the taxonomic confusion caused by the unusual *prfA*-negative phenotype. BLASTn comparison with the NCBI GenBank database yielded a definitive yet discordant outcome (Table 1).

**Table 1:**
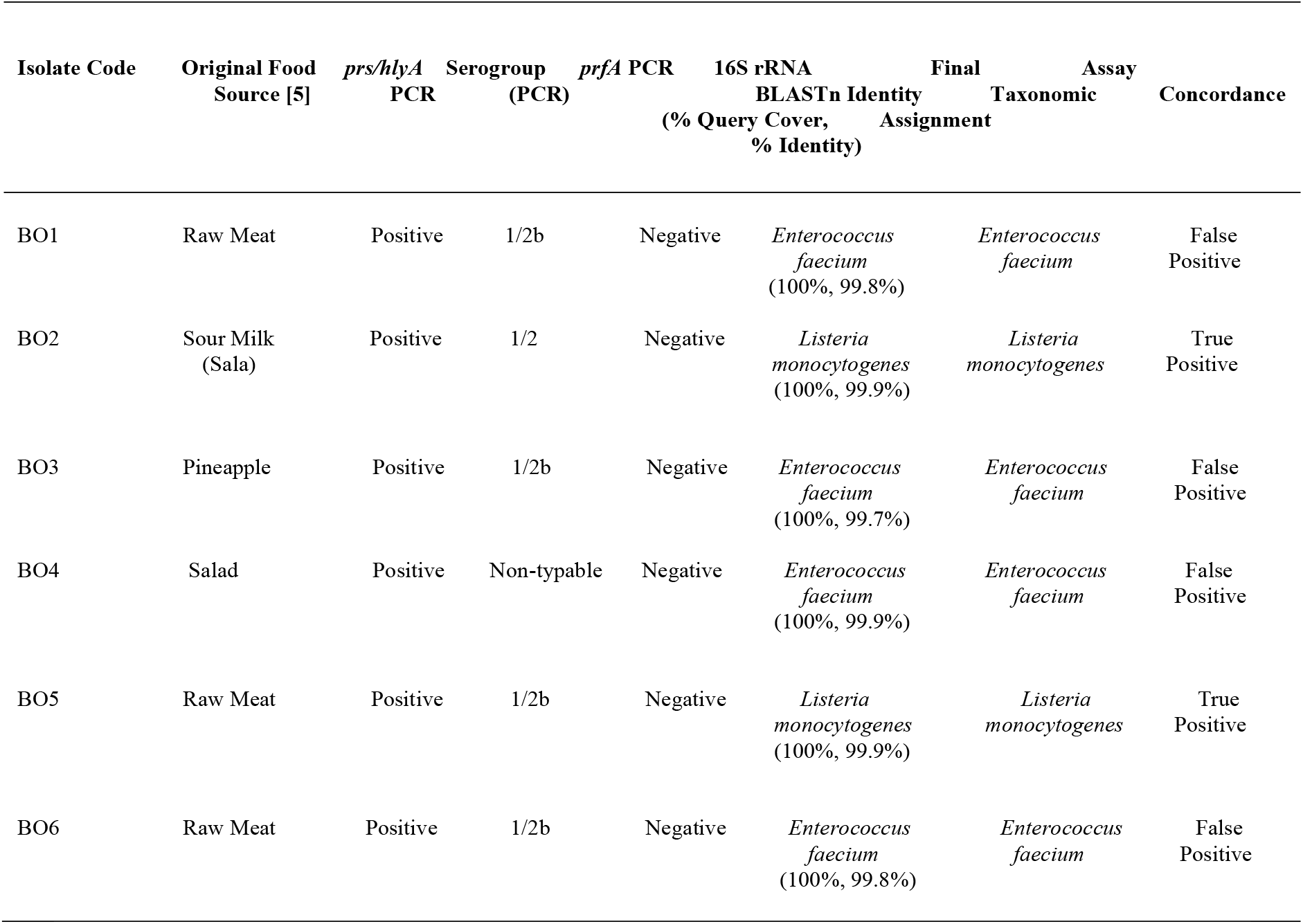
Concordance between *prs/hlyA* multiplex PCR assay and definitive 16S rRNA gene.

| Isolate Code | Original Food Source [5] | *prs/hlyA* PCR | Serogroup (PCR) | *prfA* PCR | 16S rRNA BLASTn Identity (% Query Cover, % Identity) | Final Taxonomic Assignment | Assay Concordance |
| --- | --- | --- | --- | --- | --- | --- | --- |
| BO1 | Raw Meat | Positive | 1/2b | Negative | *Enterococcus faecium* (100%, 99.8%) | *Enterococcus faecium* | False Positive |
| BO2 | Sour Milk (Sala) | Positive | 1/2 | Negative | *Listeria monocytogenes* (100%, 99.9%) | *Listeria monocytogenes* | True Positive |
| BO3 | Pineapple | Positive | 1/2b | Negative | *Enterococcus faecium* (100%, 99.7%) | *Enterococcus faecium* | False Positive |
| BO4 | Salad | Positive | Non-typable | Negative | *Enterococcus faecium* (100%, 99.9%) | *Enterococcus faecium* | False Positive |
| BO5 | Raw Meat | Positive | 1/2b | Negative | *Listeria monocytogenes* (100%, 99.9%) | *Listeria monocytogenes* | True Positive |
| BO6 | Raw Meat | Positive | 1/2b | Negative | *Enterococcus faecium* (100%, 99.8%) | *Enterococcus faecium* | False Positive |

Only two of the six PCR-positive isolates (BO2 and BO5) were identified as *L. monocytogenes* by sequencing. With 99.7–99.9% similarity, the remaining four isolates (BO1, BO3, BO4, and BO6) were determined to be *Enterococcus faecium*. In this evaluation, the *prs/hlyA* PCR assay has a false-positive rate of 66.7% (4/6).

### 3.3. Additional Analysis of PCR-Positive Isolates

Five of the six PCR-positive isolates (BO1, BO2, BO3, BO5, and BO6) belonged to serogroup 1/2b (471 bp amplicon), according to multiplex PCR for major serogroups linked to human listeriosis (Fig 1B). Using the primers, isolate BO4 was not typable. All six isolates consistently lacked the *prfA* amplicon.

### 3.4. Phylogenetic analysis uncovers intricate taxonomic relationships among isolates

Phylogenetic analysis based on partial 16S rRNA gene sequences generated a tree with surprising topological characteristics (refer to Fig. 2). Three of the four isolates initially identified as *Enterococcus faecium*—BO1, BO4, and BO6—formed a robust clade (bootstrap ≥ 90%) alongside reference strains of *E. faecium* from China (OR795718.1) and Thailand (LC027228.1), validating their classification. However, the fourth presumed *E. faecium* isolate, BO3, did not fall within this Enterococcus clade. Instead, it was phylogenetically situated between the two *Listeria monocytogenes* isolates (BO2 and BO5) within the wider *Listeria* genus clade. The two *L. monocytogenes* isolates, BO2 and BO5, reliably grouped within a *Listeria* clade that included reference strains from Germany, Poland, China, and India. They formed a unique sub-clade with moderate bootstrap support (54–57%), suggesting potential regional lineage differentiation.

**Figure 2.**
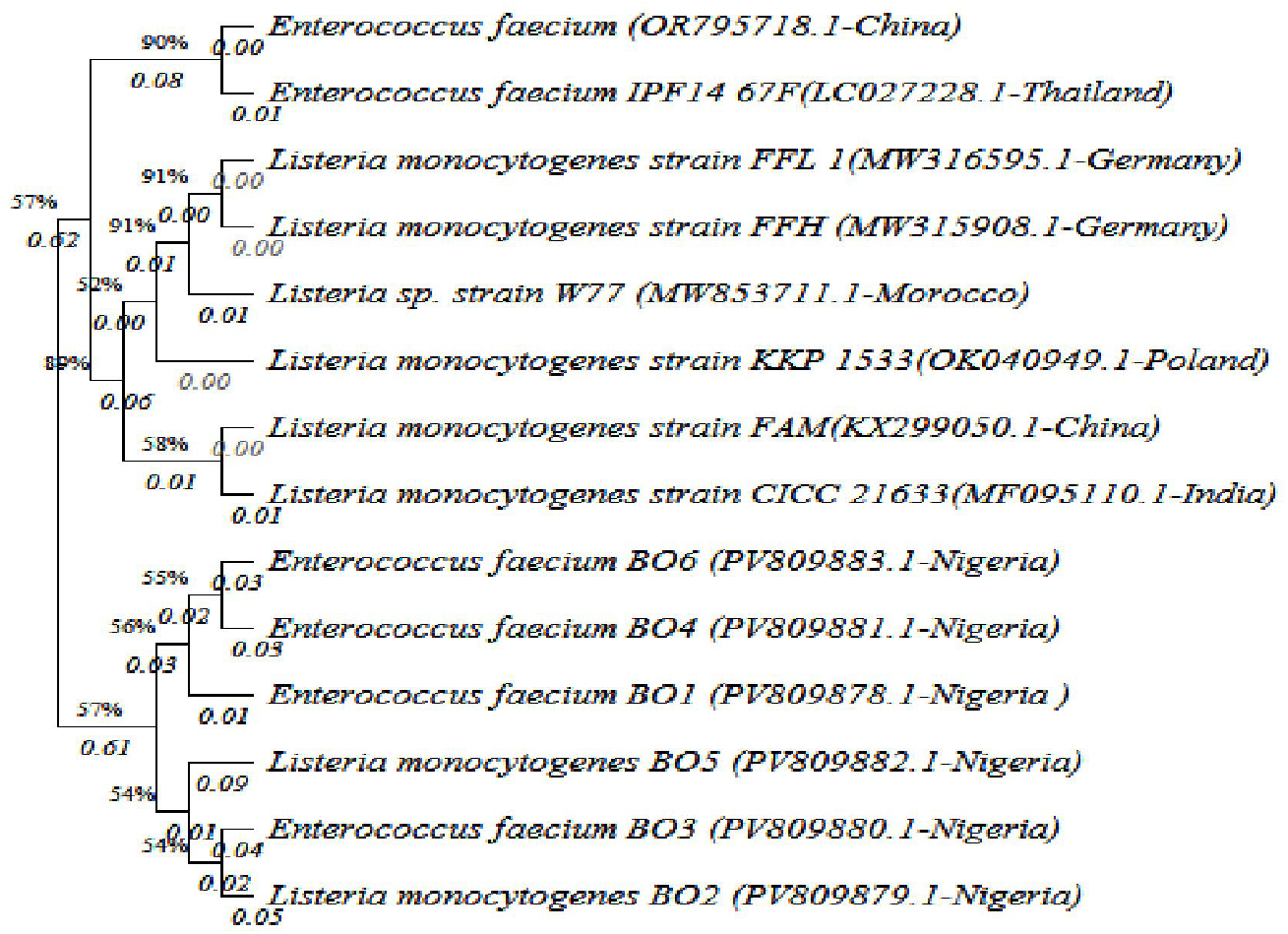
Phylogenetic analysis utilizing 16S rRNA gene sequences. A Neighbor-Joining tree illustrates the relationship between the study isolates (highlighted in bold) and reference sequences sourced from GenBank. The four *Enterococcus faecium* isolates (BO1, BO3, BO4, BO6) group within the Enterococcus clade. The two *Listeria monocytogenes* isolates (BO2, BO5) group within the *Listeria* clade. Bootstrap values (≥50%) derived from 1000 iterations are presented. The scale bar represents 0.05 nucleotide substitutions per site.

### 3.5. Overview of Assay Performance Metrics

Using sequencing as the reference standard, the performance metrics of the *prs/hlyA* multiplex PCR assay in this study were:

- Sensitivity: 100% (2 out of 2 true *L. monocytogenes* isolates detected)
- Specificity (for *E. faecium*): 0% (0 out of 4 non-target *E. faecium* isolates correctly excluded)
- False-Positive Rate: 66.7% (4 out of 6 PCR-positive results were incorrect)
- Positive Predictive Value (PPV): 33.3% (only 2 out of 6 PCR-positive results were true L. monocytogenes)

The 16S rRNA gene sequences obtained in this study have been submitted to GenBank with accession numbers PV809878.1 (BO1) to PV809883.1 (BO6).

## 4.0 DISCUSSION

This research highlights a significant specificity issue in a commonly utilized confirmatory PCR assay, illustrating that the Doumith *prs/hlyA* multiplex PCR may yield false-positive results for *Listeria monocytogenes* due to cross-reactivity with *Enterococcus faecium*. With a false-positive rate of 66.7% (4 out of 6 isolates) observed in this assessment, our results have important ramifications for diagnostic and food safety labs that depend on this assay as a conclusive test. The most likely reason for this misidentification is the binding of primers in a non-specific manner. Although the prs gene is used as a target specific to the *Listeria* genus, it encodes a highly conserved housekeeping enzyme (phosphoribosylpyrophosphate synthetase) that is found throughout bacteria, including Enterococcus [3]. The primer sequences probably display enough similarity with the relevant regions in the *E. faecium* genomes present in our sample set, resulting in successful amplification. This highlights a crucial concept in molecular diagnostics: tests that have been validated with certain strain collections from a specific geographical or ecological context may lose their specificity when used in another area with different microbial populations [3]. Additionally, recorded proof of horizontal gene transfer among Firmicutes in common environments, like food, might lead to shared genetic segments that cause cross-reactivity [14, 15]. The persistent lack of a *prfA* amplicon in all six PCR-positive isolates raised the initial and significant diagnostic concern. The *prfA* gene functions as the primary transcriptional activator of the *L. monocytogenes* pathogenicity island and is regarded as a vital, conserved element of virulent strains [12]. While there are attenuated or avirulent strains that lack a functional *prfA* gene, finding a complete absence of an amplicon in a hlyA-positive isolate is quite uncommon and should raise concerns about a potential misidentification, which was confirmed through sequencing in this instance. For the two isolates that were definitively confirmed as *L. monocytogenes* (BO2, BO5), the negative *prfA* PCR result may suggest mismatches between the primer and template due to locally varying *prfA* alleles, emphasizing a secondary limitation of the assay’s published primer set that is dependent on the strain [16].

The phylogenetic analysis based on the 16S rRNA gene offered crucial, albeit intricate, taxonomic insights that were integral to our diagnostic assessment. This analysis effectively resolved the majority of the isolates, confirming *E. faecium* isolates (BO1, BO4, BO6) and *L. monocytogenes* isolates (BO2, BO5) to their anticipated genera, but it also uncovered a notable phylogenetic inconsistency. The classification of the presumed *E. faecium* isolate BO3 within the *Listeria* clade contradicts its original phenotypic classification and highlights a key limitation of depending on single-marker phylogenetics for definitive identification in certain cases.

This discrepancy emphasizes the well-recognized limitations of the 16S rRNA gene, which, despite being regarded as a gold standard for broad taxonomic categorization and differentiation between genera like *Listeria* and Enterococcus, can produce ambiguous or misleading outcomes when sequences display unusual evolutionary paths. For BO3, possible explanations could be 1) an initial misidentification, 2) the acquisition of a *Listeria*-like 16S rRNA gene through horizontal gene transfer, or 3) the occurrence of a chimeric PCR artifact. Regardless of the reason, this observation prompted the assay’s failure to accurately identify this isolate, as the molecular target diverged from the anticipated Enterococcus sequence.

The grouping of the two confirmed *L. monocytogenes* isolates (BO2, BO5) into a distinct sub-clade, separate from global reference strains, implies potential local genetic divergence that aligns with the formation of region-specific ecotypes. However, the primary significance of this phylogenetic analysis was its function as a high-resolution comparator. It not only confirmed the assay’s effectiveness for isolates with clear 16S rRNA phylogeny but, more crucially, uncovered its diagnostic failure by indicating a major taxonomic inconsistency that simpler comparative methods might overlook.

The practical ramifications for laboratory operations are considerable. Laboratories that rely solely on the *prs/hlyA* multiplex PCR as a confirmatory test risk producing inaccurate results, which could lead to unnecessary product recalls, inflated epidemiological statistics, and misallocation of public health resources. Thus, we recommend a revised diagnostic protocol in which positive results from this assay are treated as presumptive. It is essential to confirm findings through an orthogonal, sequence-based method. For resource-limited environments, sequencing the 16S rRNA gene of a representative subset of amplicons presents a cost-effective approach for periodic assay validation. Where possible, adding a second highly specific target (such as *iap*) into a confirmatory PCR is advisable to improve reliability.

In conclusion, this case study underscores the importance of context-specific validation for even established molecular assays. It demonstrates how an anomalous result (*prfA*-negativity) can act as a crucial indicator of a more profound diagnostic issue. By combining targeted PCR with definitive sequencing, laboratories can greatly enhance the accuracy of *L. monocytogenes* reporting, thus reinforcing food safety measures and public health diagnostics.

## ACKNOWLEGEMENT

We thank the Department of Medical Microbiology University of Maiduguri Teaching Hospital, Department of Veterinary Public Health, Ahmadu Bello University Zaria, and the Department of Microbiology, University of Nigeria Nsukka, for their technical support.

## FUNDING

This study did not receive any specific funding from public, commercial, or not-for-profit organizations.

## CONFLICT OF INTEREST

The authors state that there are no conflicts of interest.

## AUTHOR CONTRIBUTIONS

H.B.A carried out the conceptualization, methodology, investigation and wrote the original draft. A.S.K supervised, validated and reviewed the methodology. A.J.M., Y.U., A.S.B. and M.U carried out investigation, validation and reviewed the write up. All authors approved the final manuscript.

## DATA AVAILABILITY

The sequences of the 16S rRNA gene produced in this study have been submitted to the NCBI GenBank database with accession numbers PV809878.1 to PV809883.1.

## REFERENCES

1. Gandhi M, Chikindas ML. Listeria: A foodborne pathogen that knows how to survive. Int J Food Microbiol. 2007;113(1):1–15.

2. Ryu J, Park SH, Yeom YS, Shrivastav A, Lee SH, Kim YR, et al. Simultaneous detection of Listeria species and Listeria monocytogenes using a multiplex PCR assay. J Microbiol Biotechnol. 2013;23(1):90–9.

3. Doumith M, Buchrieser C, Glaser P, Jacquet C, Martin P. Differentiation of the major Listeria monocytogenes serovars by multiplex PCR. J Clin Microbiol. 2004;42(8):3819–22.

4. Rantsiou K, Ferranti P, Cocolin L. Limitations of 16S rRNA gene sequencing for species-level identification of foodborne bacteria. Curr Opin Food Sci. 2021;42:173–9.

5. BA Haruna, Chiwar HM, Kumurya AS, and H.J Balla. Assessment of Antimicrobial Resistance Trends in Listeria monocytogenes in Borno State: Implications for Food Safety and Public Health. UMYU Journal of Microbiology Research. 2025;10(2):1–10.

6. Liu D, Lawrence ML, Ainsworth AJ, Austin FW. Comparative assessment of acid, alkali and salt tolerance in Listeria monocytogenes virulent and avirulent strains. FEMS Microbiol Lett. 2005;243(1):373–8.

7. Weisburg WG, Barns SM, Pelletier DA, Lane DJ. 16S ribosomal DNA amplification for phylogenetic study. J Bacteriol. 1991;173(2):697–703.

8. Tamura K, Stecher G, Kumar S. MEGA11: Molecular Evolutionary Genetics Analysis Version 11. Mol Biol Evol. 2021;38(7):3022–7.

9. Saitou N, Nei M. The neighbor-joining method: a new method for reconstructing phylogenetic trees. Mol Biol Evol. 1987;4(4):406–25.

10. Felsenstein J. Confidence limits on phylogenies: An approach using the bootstrap. Evolution. 1985;39(4):783–91.

11. Tamura K, Nei M, Kumar S. Prospects for inferring very large phylogenies by using the neighbor-joining method. Proc Natl Acad Sci U S A. 2004;101(30):11030–5.

12. Vázquez-Boland JA, Kuhn M, Berche P, Chakraborty T, Domínguez-Bernal G, Goebel W, et al. Listeria pathogenesis and molecular virulence determinants. Clin Microbiol Rev. 2001;14(3):584–640.

13. Johnson J, Jinneman K, Stelma G, Smith BG, Lye D, Messer J, et al. Natural atypical Listeria innocua strains with Listeria monocytogenes pathogenicity island 1 genes. Appl Environ Microbiol. 2004;70(7):4256–66.

14. Bertsch D, Rau J, Eugster MR, Haug MC, Lawson PA, Lacroix C, et al. Comparative genomics of Listeria species reveals a conserved genomic organization but diversity in virulence gene clusters. BMC Genomics. 2014;15:855.

15. Komora N, Maciel C, Ferreira V, Martins M, Castro P, Teixeira P. The wall teichoic acid polymerase TagF is a new target for Listeria phage and Enterococcus phage. Sci Rep. 2020;10:12151.

16. Painset A, Björkman JT, Kiil K, Guillier L, Mariet JF, Félix B, et al. LiSEQ – whole-genome sequencing of a cross-sectional survey of Listeria monocytogenes in ready-to-eat foods and human clinical cases in Europe. Microb Genom. 2020;6(2):e000257.

17. Doumith M, Buchrieser C, Glaser P, et al. Differentiation of the major Listeria monocytogenes serovars by multiplex PCR. J Clin Microbiol. 2004;42(8):3819–3822.

18. Swetha CS, Suneetha P, Harshini M. Molecular detection of virulence genes in Listeria monocytogenes isolated from dairy products. J Appl Nat Sci. 2012;4(1):75–79.

19. Notermans S, Dufrenne J, Leimeister-Wächter M, et al. Use of a tissue culture assay to detect invasive Listeria monocytogenes among wild-type strains isolated from food and humans. J Appl Bacteriol. 1991;70(2):121–126.

